# The genetic code assembles via division and fusion, basic cellular events

**DOI:** 10.1101/2023.05.01.538992

**Authors:** Michael Yarus

**Affiliations:** Department of Molecular, Cellular and Developmental Biology, University of Colorado Boulder, Boulder, CO 80309-0347

## Abstract

Standard Genetic Code (SGC) evolution is quantitatively modeled in computed ‘worlds’ containing up to 2000 independent coding ‘environments’. Environments can host multiple codes that may fuse or divide, with division yielding identical descendants. Code division may be selected - sophisticated gene products could be required for orderly separation. Several unforeseen results emerge: more rapid evolution requires unselective code division, rather than its selective form. Combining selective and unselective code division, with/without code fusion, with/without independent environmental coding tables and with/without wobble defines 2^5^ = 32 possible pathways for SGC evolution. These 32 possible histories are compared, particularly, for speed and accuracy. Pathways differ greatly; for example, ≈ 300-fold different in time to evolve SGC-like codes. Eight of 32 pathways, employing code division, are quickest. Four of these eight, that combine fusion and division, also unite speed and accuracy. The two precise, swiftest paths, thus the most likely routes to the SGC, are similar, differing only in fusion with independent environmental codes. Code division instead of fusion with unrelated codes implies that independent codes can be dispensable. Instead, a single ancestral code that divides and fuses can initiate fully encoded peptide biosynthesis. Division and fusion create a ‘crescendo of competent coding’, facilitating search for the SGC, and also assist advent of otherwise disfavored wobble coding. Code fusion readily unites multiple codon assignment mechanisms. But via code division and fusion, the SGC is shown to emerge from a single primary origin, via familiar cellular events.

## Introduction

### The problem

Automated total searches of ≈ 2.5 x 10^5^ bacterial and archaeal genomes (Shulgina and Eddy 2021, 2023) find only slightly altered genetic codes, related to the Standard Genetic Code (SGC). Hence, true alternative codes are exceedingly rare on the modern Earth. Modern biota therefore convincingly trace to a single ancestral group encoding peptides using a close SGC relative. This ancient all-inclusive ancestor presents a problem of ultimate significance for Biology, and is the topic here.

### Incorporating code division

We follow the evolution of numerous coding tables through time in code evolution environments, using Monte Carlo kinetics (Yarus 2021; see Methods). The time for one round of evolution in a code environment is called a passage. An evolving environment undergoing a passage containing zero, one, tens or hundreds of coding tables can <u>either</u>: initiate coding (probability = Pinit) in a first table with a random assignment, add a new table (Ptab), beginning by assigning a random codon to a randomly chosen function (20 amino acids, start, stop). Or finally, an environmental code can divide (Pdiv), adding an additional code identical to the pre-existing one. Code division can be non-selective, allowing any code to divide, or a **<u>c</u>**ompleteness **<u>c</u>**riterion (cc) can specify that division occurs only after a certain variety of coding assignments are possible. A completeness criterion recognizes that specific folding (Solis 2019) and enzymatic function (Akanuma et al. 2002; Shibue et al. 2018; Kimura and Akanuma 2020) is observed only when proteins contain 10-13 amino acids; a proteome capable of precise division plausibly requires prior code evolution to encode multiple amino acids.

An experiment may comprise many contemporaneous environments (a world), which reports its status when its environments reach a set goal, like SGC-level completeness (e.g., encoding ≥ 20 functions) or at a specified time (e.g., after 121 passages).

## Results

### Presenting effects of code evolution

Worlds composed of perhaps hundreds of environments with variably evolved codes present problems of exposition. Multiple presentation problems can be solved by the method in Fig. 1A. Mean data is listed in a specific numbered order (#1, 2, 3…) reflecting different underlying mechanisms; thus mechanism appears as periodicity in the plot. Behavior in multiple mechanistic dimensions can be read from an ordinary two-dimensional graph.

**Figure 1A.**
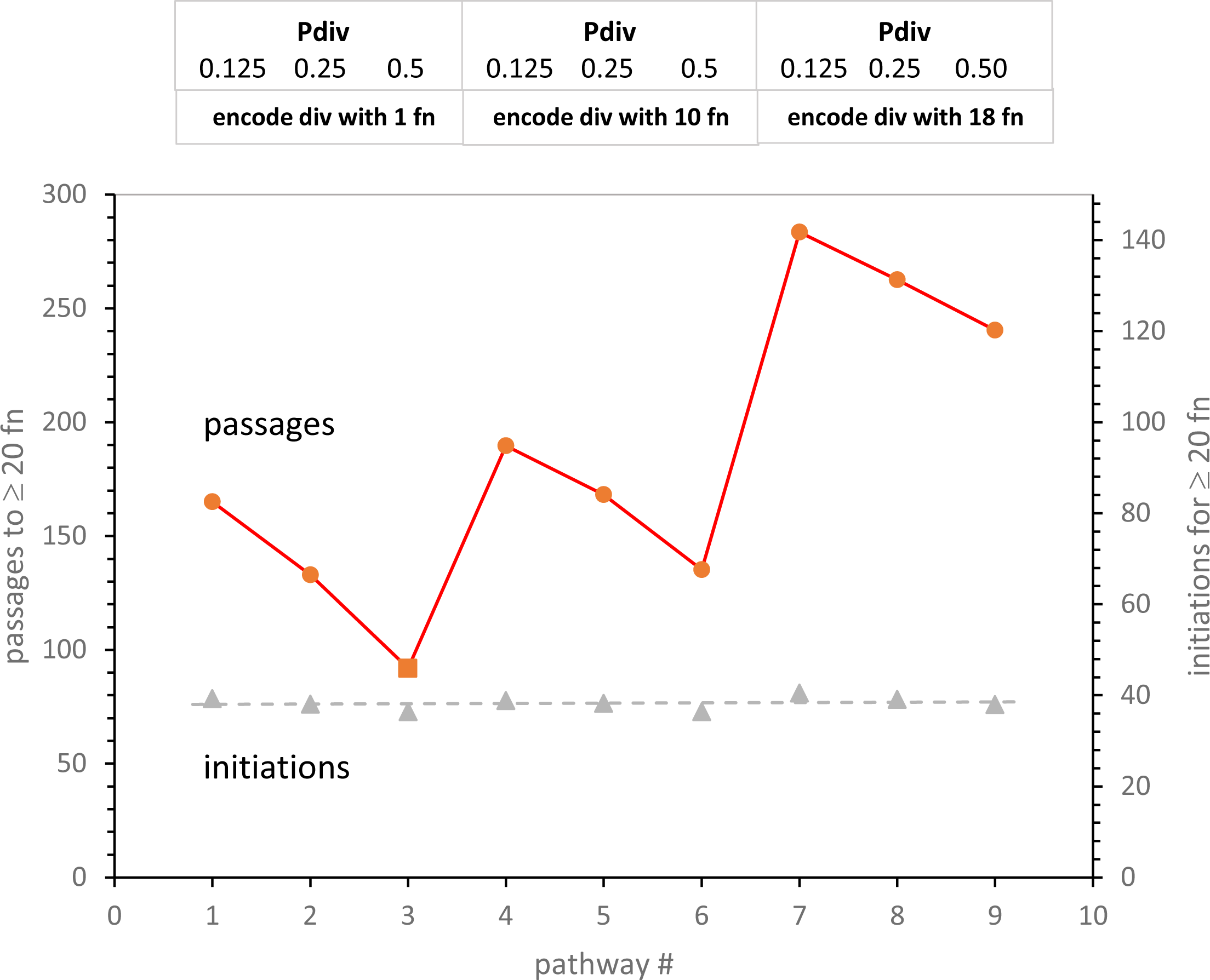
Effects of code division on time to evolve ≥ 20 encoded functions, and on number of initial assignments required for ≥ 20 encoded functions. The pathway x-axis follows a structured list of code division variables (see text) named in the box at graph top: “Pdiv” = probability of unselected code division/passage, “encode div with..” = code completeness (cc) required to encode accurate code division. Pmut=0.00975, Pdec=0.00975, Pinit=0.15, Prand=0.05, Pfus=0.001, Ptab=0.08, Pwob=0.0 – results show means for evolution in 500 environments.

Structured plotting is illustrated (Fig. 1A) by three groups of codes that have a different threshold for code division (cc, the **<u>c</u>**ompleteness **<u>c</u>**riterion) at 1, 10 and 18 codon assignments. Within each such group, codes have an unselective probability of division per passage of 0.125, 0.25 and 0.5. Thus one can read effects of increasing division within groups of three, and also read the effects of different division thresholds by comparing such groups.

### Effects of code division

In Fig. 1A, increased division (Pdiv) always reduces time to evolve SGC-like coding, ≥ 20 assigned functions. Quicker evolution is slightly less for the same Pdiv change at higher threshold (comparing mean slopes of threes). In addition, evolution is increasingly rapid if the division threshold is lowered from near-completion (set at ≥ 20 functions encoded) to no threshold at all on the left (threshold at 1 function; any code can divide). Fig. 1A presents a non-trivial result: non-selective code division (mechanism #3, red square), acting throughout evolution, evoves SGC-like coding most quickly.

### Division effects on rate

Division is revisited in Fig. 1B, plotting speed of evolution versus number of code divisions to reach SGC-like assignment. In Fig. 1B, time to ≥ 20 encoded functions with Fig. 1A’s variety of division probabilities and thresholds, declines rapidly as code divisions increase. Fastest mean SGC-like evolution with code division, 92 passages, is more than 3 times faster than previously with these same code passage probabilities (Yarus 2022a).

**Figure 1B.**
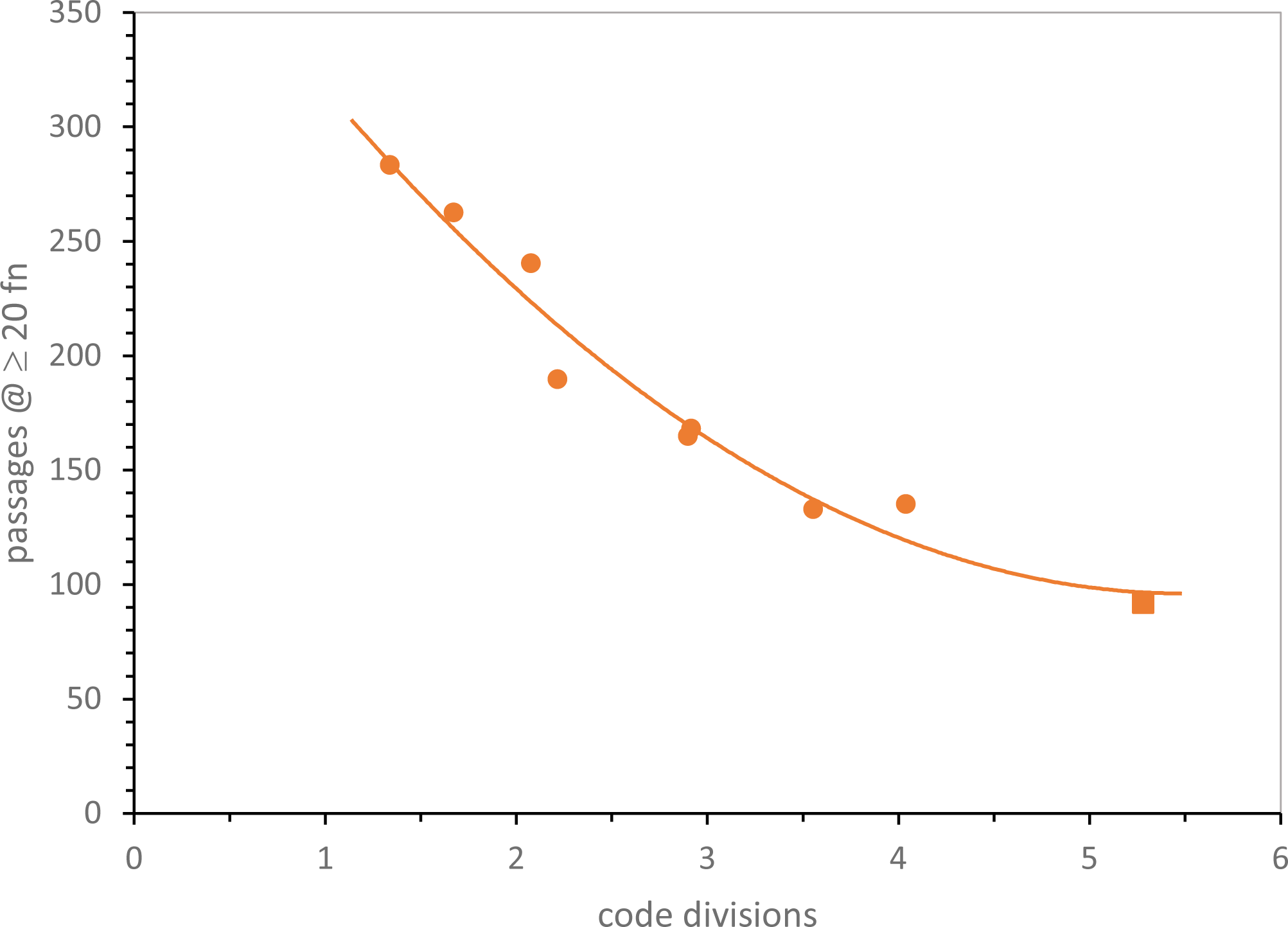
Mean time to evolve ≥ 20 encoded functions versus mean number of code divisions for those codes. A square marks shortest evolutionary time. Environments are those in Fig. 1A.

### Small effects

A mechanism-structured plot is also useful when substantial effects are absent. Fig. 1A plots the number (≈38) of random initial codon assignments required to reach ≥ 20 assigned functions, on its rightward axis. This is hardly altered in mechanisms 1 through 9. Close inspection of displacements from the least-squares dashed line discloses periodic behavior; fast evolution requires slightly fewer assignments. But the structured plot (Fig. 1A) highlights how small this effect is for these changes in code division. Fixed assignments are not a rule; pathways <u>can</u> use assignments inefficiently. But even conditional constancy will be useful below, to clarify complex evolution.

### Presenting code accuracy

A general measure of SGC similarity is frequently needed. One would like to avoid assigning many codons, but to different functions than in the historical SGC.

In this work, misassigments (mis) with respect to the biological SGC are counted (Yarus 2021d). Codes with no difference from SGC assignment are denoted mis0, those with one difference are mis1 codes, and on to mis2, mis3… The fraction of SGC-like assignments provides an index of distance that meets our need to measure evolutionary accuracy.

However, this can pose a problem of precision: SGC-identical mis0 codes can be infrequent, even pragmatically unmeasurable for inaccurate evolutionary modes. However, Fig. 2 shows how this problem can be met. The distribution of errors is smooth and unimodal – the fraction of SGC-like codes (here, the fraction that are mis0 and mis 1) rises smoothly with the decrease of mean mis in near-complete codes. Because mean mis are measured in up to two thousand environments, average misassignment is always known with significant precision. Proximity to the SGC is therefore measured (Fig. 2) either by calculating mean misassignment (mis; accuracy better when smaller) among most complete codes, or by counting codes nearest the SGC when accessible (accuracy better when larger).

**Figure 2.**
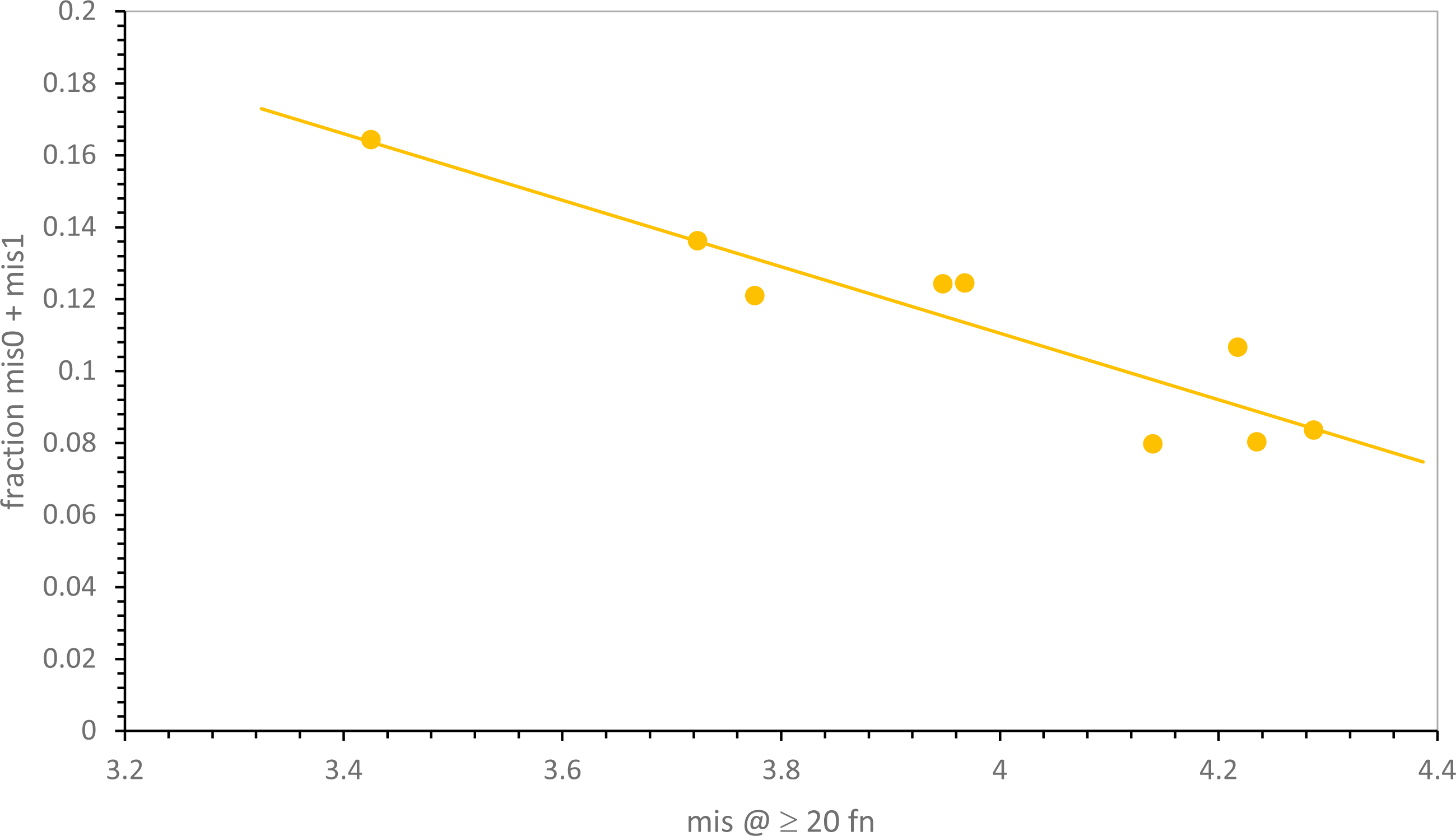
Mean mis and mean fraction of codes with SGC-like assignments are closely, and inversely, related. Mis0 = identical to SGC assignments; mis1 = one difference from SGC assignments. Environments are those in Fig. 1A.

### Mechanisms and code accuracy

Accuracy as mean misassignments in Fig. 3, like time in Fig. 1A, is plotted versus Fig. 1A’s division pathways 1 to 9. A Fig. 1A-like pattern reappears. So: code accuracy is greatest with more division (greater Pdiv) in several contexts. The sensitivity of code accuracy to division frequency declines significantly as a division threshold increases (Fig. 3). Absolute accuracy also is greatest when the threshold (completeness criterion, cc, square marker) is low: pathway #3 is most accurate (Fig. 3). Most accurate code evolution utilizes frequent division, and approaches the SGC quickly without selection for coding sophistication; any code at all meets a one-assignment division “threshold”. This result reappears in a much more complex mechanistic context below.

**Figure 3.**
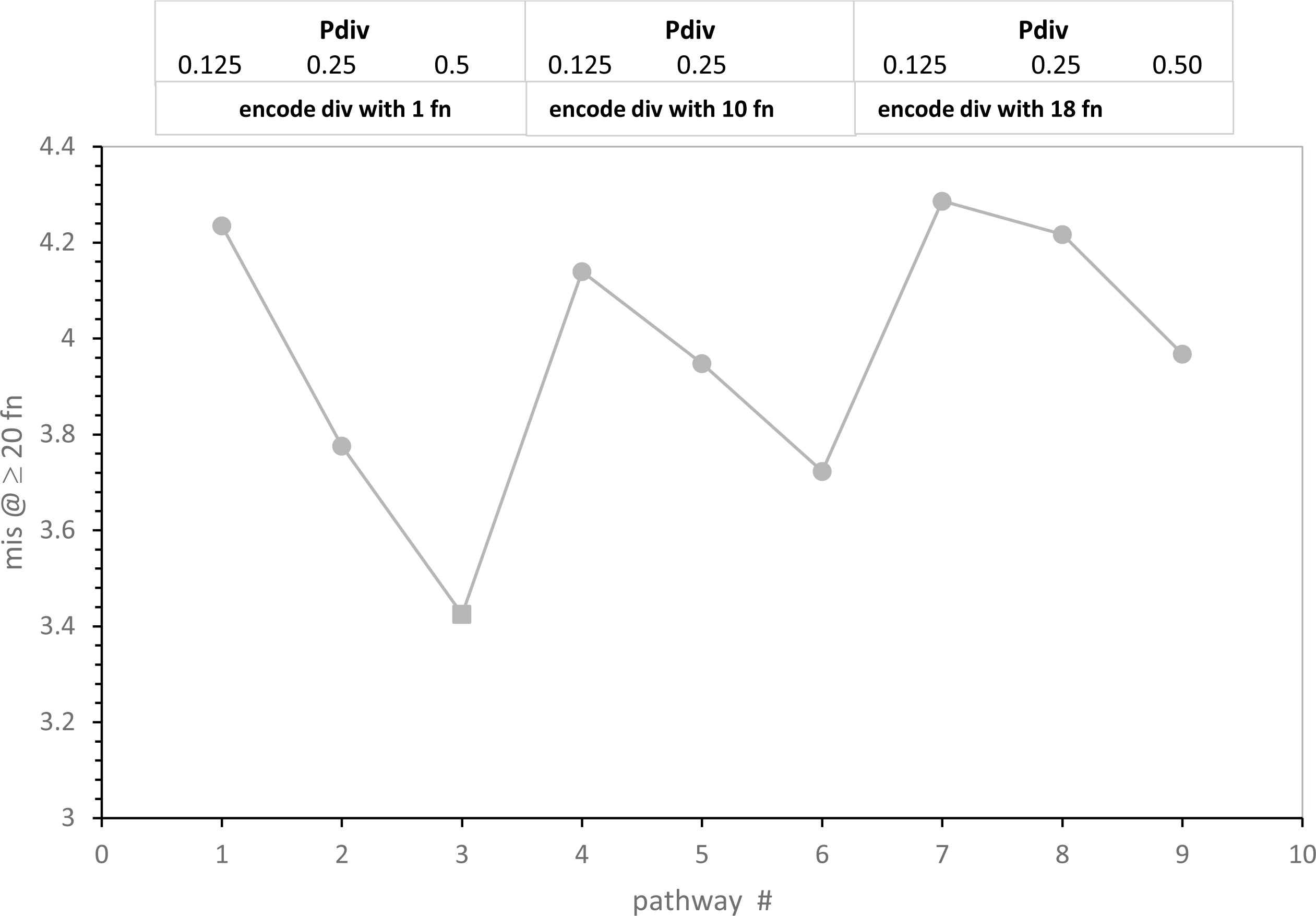
Effects of division variables (Pdiv and cc) on accuracy of evolution to ≥ 20 encoded functions. The x-axis is a structured list of pathways like Fig. 1A. Environments are those in Fig. 1A. A square marks the most accurate pathway.

Moreover, in Fig. 3, code division has an interesting property previously shown for code fusion (Yarus 2022a): more division (greater Pdiv) reduces error, implying constraint of the present mixture of initial SGC and random assignments. Such adherence to an underlying coding consensus (Fig. 3) is weakened if a threshold delays initiation of code division. But it is more division, not division selecting code progress, that produces an accurate code (Fig. 3) while also evolving it quickly (Fig. 1A).

### Five-dimensional comparison of 32 pathways

Incorporation of division effects into a Monte Carlo kinetic scheme (Methods) for code table evolution defines 32 pathways toward the genetic code; with/without code division (probability of division, as well as division threshold), with/without code fusion (Yarus 2022a), with/without independent coding tables (Yarus 2022a), with/without simplified Crick wobble (Yarus 2021c). These pathways are quantitatively compared in Fig. 4, using the structured display method of Fig. 1A to organize five-dimensional data (see the supplementary file).

**Figure 4.**
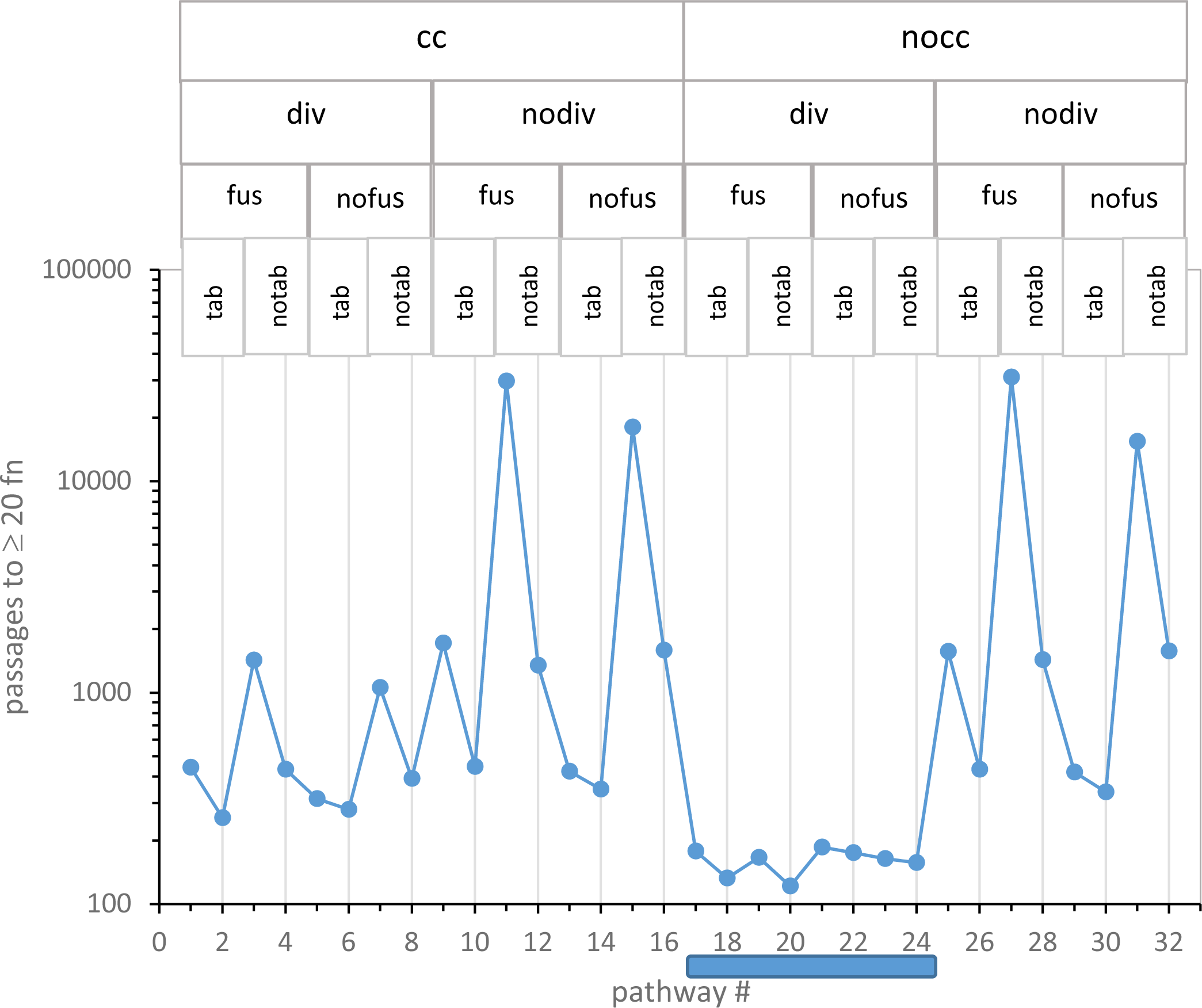
Logarithmic mean time to evolve ≥ 20 encoded functions for 32 code evolution pathways. Pathway mechanism abbreviations are listed at graph top: cc = require <u>c</u>ompleteness <u>c</u>riterion for code division, nocc = no cc required; div = allow code division with probability Pdiv, nodiv = no code division; fus = allow codes to fuse with probability Pfus, nofus = no code fusion; tab = allow independent environmental coding tables, origin probability Ptab; notab, no parallel tables; wob = allow wobble coding (no vertical line through point), nowob = simple base pairing (vertical line); A shaded bar below the x-axis marks the nocc div canyon. Numerical data are in a supplementary file.

Fig. 4 presents the time to reach ≥ 20 encoded functions (in passages, ordinate) versus all 32 numbered mechanisms on the x-axis. For example, minimal time to evolve ≥ 20 encoded functions occurs via mechanism #20, which (reading titles above and the vertical line through the point: legend, Fig. 4) utilizes no completeness threshold for division (nocc), incorporates probable code division (div), allows codes to fuse (fus) but has no independent codes forming in its environment (notab), and evolves during initial assignments in the absence of wobble (nowob vertical line). Path #20 reappears frequently below.

### A glance identifies the fastest evolution

In Fig. 4, mechanisms that have no completeness threshold (nocc: cc = 1) and probable code division (div) form a “canyon” (mech #17-24; shaded bar) each of whose 8 pathways evolve ≥ 20 encoded functions faster than <u>any</u> of the other 24 mechanisms examined.

Moreover, this nocc div canyon is the major difference between Fig. 4 left and right. Superior unselective code division, first seen in Fig. 1A, reappears here in a broader mechanistic context. Therefore the path of least selection, that is, the probable evolutionary path, will be a nocc div unselective route. Accordingly, code division greatly changes early code evolution, and nocc div will be required elements in best SGC pathways.

### A glance identifies the slowest evolution

In Fig. 4, the four slowest routes to the SGC have in common that codes do not divide (nodiv) and no additional codes appear alongside from independent code origins (notab). Under such conditions, fusion is irrelevant because there are no additional codes to fuse. Thus for these four slowest pathways, fus/nofus mechanisms are about equivalently poor, because code fusion is inaccessible and irrelevant. A single code in each environment must evolve alone to SGC proximity, and this requires a complex set of events, with many digressions, making these the most improbable evolutionary routes. This matches previous observations (Yarus 2022a), and rationalizes emphasis on superior pathways below, all of which exploit code-code interactions.

### Wobble is always inhibitory

Among 32 pathways in Fig. 4, 16 encode using wobble, and 16 do not. One can consider the 16 wob (no vertical line)/nowob (line) pairs together by noting that each wobbling pathway (no vertical line) is accompanied by a non-wobbling pathway immediately to its right (line) that differs only in lacking simple Crick wobble (Yarus 2021b).

Mechanisms differ in their sensitivity to wobble. Slow single-coding-table environments are very much impaired if they must use wobble assignments. In contrast, the 8 mechanisms of the nocc div canyon (Fig. 4) are less sensitive to inhibitory wobble effects. However, throughout all 16 wob/nowob pairs in Fig. 4, wobble prolongs evolution to the SGC. This extends previous findings that assignments that commit more triplets always impede progress toward complete coding (Yarus 2021c, 2021a), and that wobble specifically disrupts the evolution of codes that most resemble the SGC (Yarus 2023). Here wobble is negative in 16 varied mechanistic contexts. This strongly reinforces previous structural arguments; accurate wobble requires a complex ribosomal isomerization (Ogle et al. 2001; Yarus 2022a) and a complex tRNA structure (Schultz and Yarus 1994; Shepotinovskaya and Uhlenbeck 2013) – thus wobble encoding appeared late in code evolution, after most functions were assigned.

Addition of simple Crick wobble to present codes adds, minimally, two misassignments because unique SGC encodings, AUG/Met and UGG/Trp, are not accounted for here. Unique assignments are most simply explained as survivors from the early non-wobbling era defined just above. However, the crucial code transition from unique to wobbling assignments can definitely bear more thought.

### Independently-originating codes (tab) can speed SGC evolution, but not in the nocc div canyon

The effect of multiple independent codes arising side-by-side, then interacting within an SGC-evolving environment, can also be assessed in Fig. 4. Pairs of tab/notab mechanisms, in which the only change is absence of independently-evolving coding tables, have sequential odd or even numbers.

For example, mechanisms #10 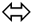 #12 and #18 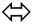 #20 differ only in lacking parallel environmental codes in higher-numbered mechanisms. But these two pairs differ greatly in the resulting effect. Loss of other codes slows SGC evolution significantly on the left in Fig. 4 (#10 to 12; 447 to 1345 passages), where nodiv cuts off other codes arising by division. In contrast, on the right (#18 to 20; 133 to 121 passages), with a supply of alternative fusion partners available from code division, parallel independent codes are instead slightly *inhibitory* to evolutionary progress. Similarly for all codes: in 12 tab/notab pairs outside the canyon, and each of 4 such pairs within the #17 to 24 canyon, codes arising by code division are always more favorable partners than independent coding tables. The gathering of coding information from several, into one, nascent code returns in Discussion.

### Speed and accuracy are related

Given that genetic codes can adhere to underlying consensus assignments (Yarus 2022a), the existence of such adherence (as in Fig. 3), as well as evolutionary speed (as in Fig. 4), is of importance. For the highly varied 32 possible mechanisms, as for the smaller, more uniform group of code divisions (Fig. 2), the fraction of codes near the SGC increases as the mean number of misassignments declines. That is, the distribution of error sharpens regularly as mean misasignment in ≥ 20 function codes declines, drawing in toward an SGC consensus. In Fig. 5, paralleling Fig. 2 for division variation only, mean misassignment (mis) is a useful measure of SGC proximity, represented as the sum of mis0 and mis1 code fractions. In fact, SGC-like codes increase more rapidly as mean misassignment closes in on the SGC, yielding sensitive detection of SGC proximity (Fig. 5).

**Figure 5.**
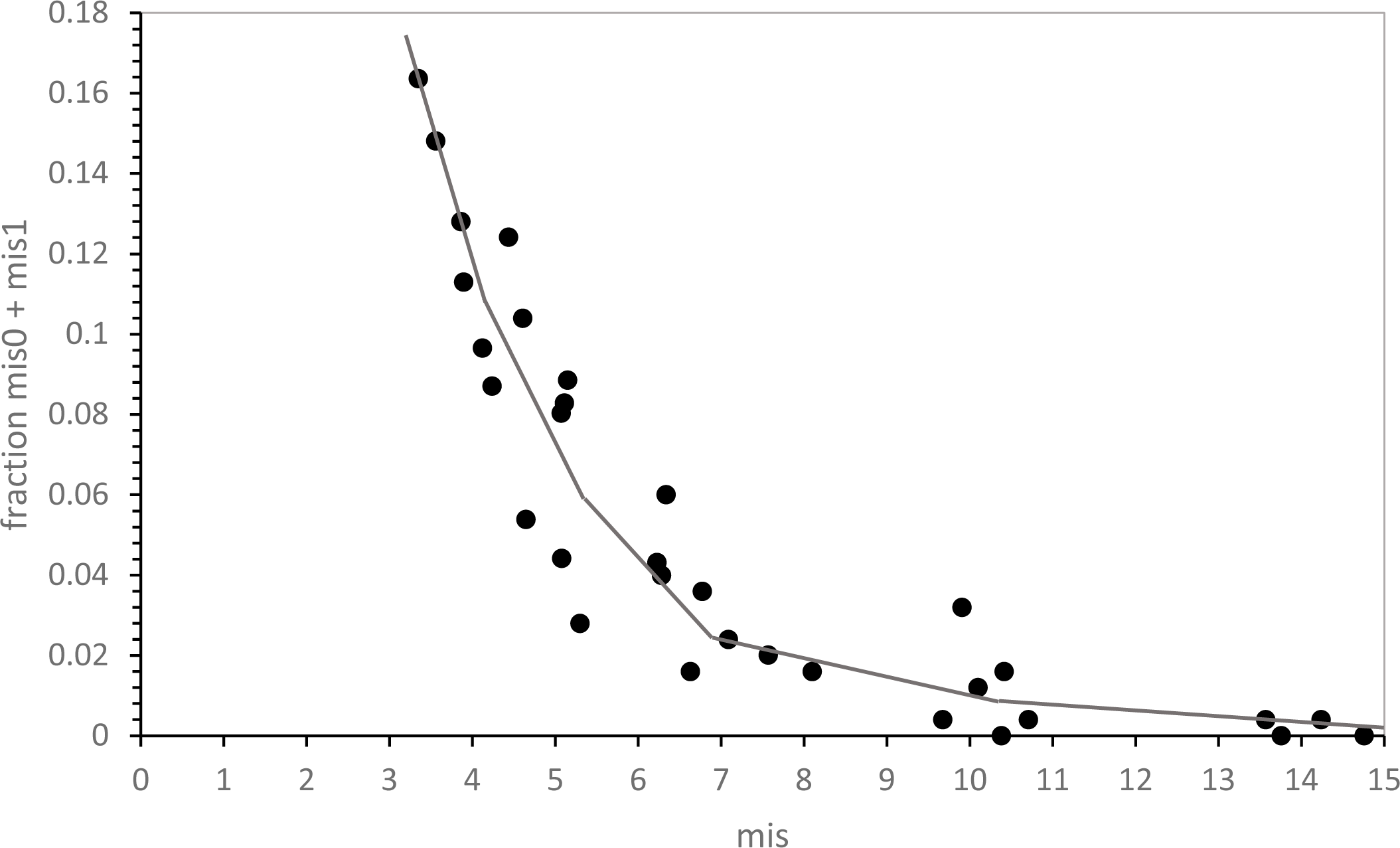
SGC-like encoding (mis0 and mis1) versus mean mis at ≥ 20 encoded functions. Environments are those in Fig. 4.

In Fig. 6, mis is plotted for the complete structured set of 32 pathways. Comparison of Figs 4 and 6 shows that evolutionary speed and accuracy are related; the two plots are similar over most pathways. For example, there is again a mechanism #17-24 nocc-div canyon, within which the lowest global error appears. However, small differences in speed and accuracy from independent tables are observed (e.g., pathway #9).

**Figure 6.**
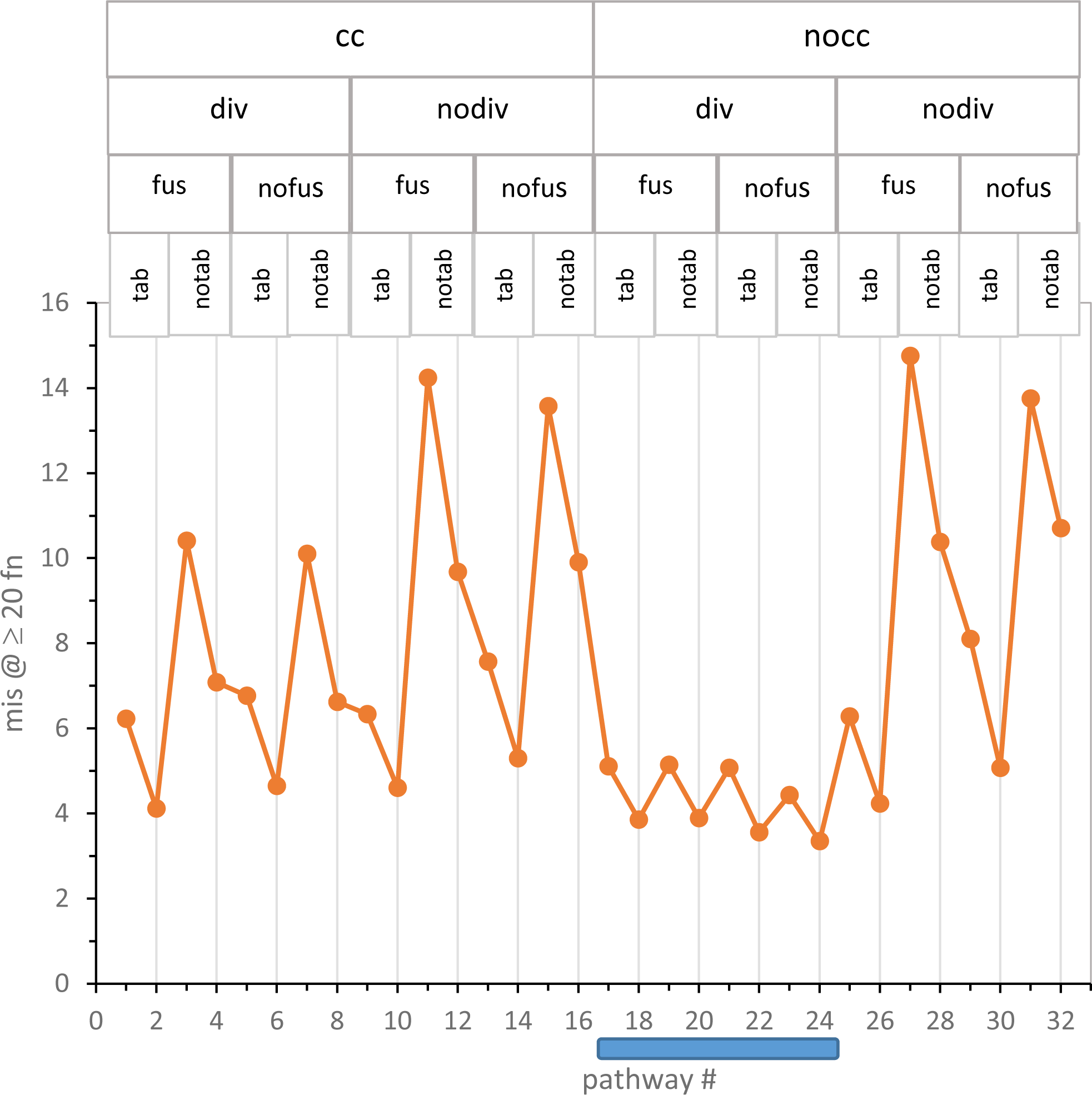
Mean misassignment (mis) at ≥ 20 encoded functions for 32 code pathways. Pathway mechanism abbreviations are listed at graph top. Environments are those in Fig. 4. The shaded bar beneath the x-axis marks the nocc div canyon.

Fig. 7 makes explicit this interaction between speed and accuracy by plotting time to ≥ 20 encoded functions in passages vs mis. There is a clear relation, though with some variation: the least squares line accounts for 86% of variance in misassignment. Fast evolution tends to occur using pathways that also approach SGC consensus. Fig. 1A, 3, 4 and 6 convey a decisive property of code evolution: it is not necessary to choose between rapid code evolution and code adherence. There are quick routes to SGC-like codes that also adhere to an underlying SGC-like consensus.

**Figure 7.**
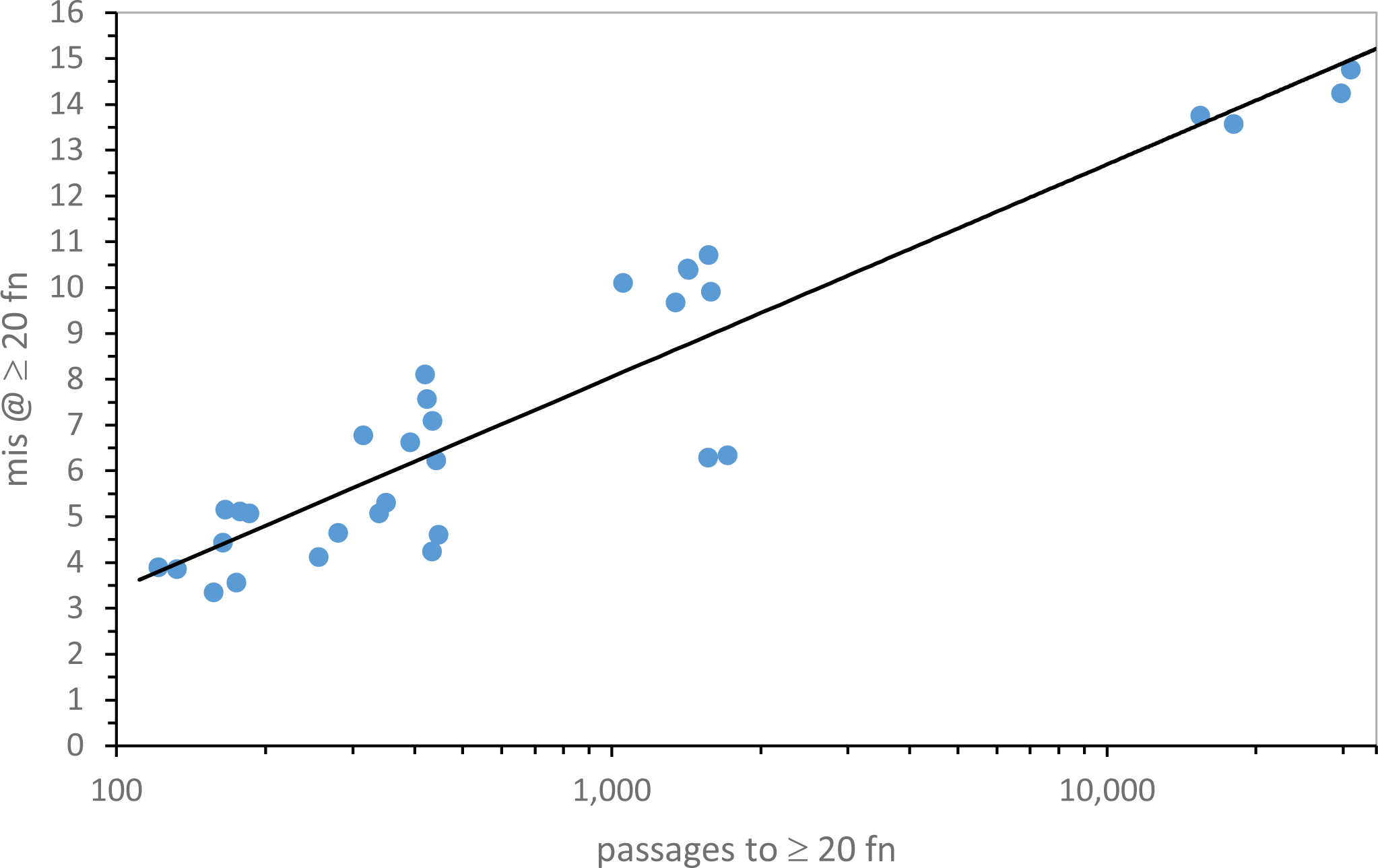
Mean misassignment at ≥ 20 encoded functions versus time for its evolution in passages, for 32 code evolution pathways. Environments are those in Fig. 4.

More quantitatively, mechanisms #18 and 20 most quickly present near–complete codes (Fig. 4). These pathways have low levels of missassignment: more than a quarter of all ≥ 20 function codes are 0, 1 or 2 assignments from the SGC. In fact, codes identical to the SGC (mis0) are more than 1 in 40 of these near-complete coding tables. Nocc-div canyon codes again provide a least selection, that is, an evolutionary route favored because it requires least selected alteration to become the SGC (Yarus 2022b).

### Distinguishing canyon codes

To focus discussion, mechanisms #18 and 20 are put foremost because they most rapidly produce complete coding (Fig. 4). As Fig. 6 shows, they do not precisely correspond to maximal resemblance to the SGC; canyon pathways 22 and 24 have slightly greater mean SGC similarity.

Differences between canyon codes appear small, but are significant. Between ≥ 20 functions in mechanisms #20 and #18, 11.2 passages intervene. Given their standard errors in 1000 environments each, a two-tailed, unequal variance t-test yields 1.8 x 10^-15^ as probability that these mean times are the same. Thus, time profiles in Fig. 4 convey statistically valid differences. Pathway #20 really arrives at ≥ 20 encoded functions before #18. However, this leaves open an important question.

### What code differences are significant?

Are Fig. 4’s time differences, however statistically significant, of importance to evolution? This question can be approached quantitatively, using the notion of least selection (Yarus 2022b). Fig. 8 combines code completeness and accuracy in one metric. The abundance of codes that both encode ≥ 20 functions (completeness) and also are accurate (fewest differences from SGC assignments) is taken as the distance to be crossed by selection. This is most relevant at early times, when such codes are first exposed to selection. In Fig. 8, the mean time to encode ≥ 20 functions for mechanism #20, 121 passages (Fig. 4), is taken as reference. Fig. 8 plots SGC proximity for all 8 canyon-bottom mechanisms at that time, using the same structured list as Fig. 4 and Fig. 6. Relevant pathway abbreviations again appear above each datum.

**Figure 8.**
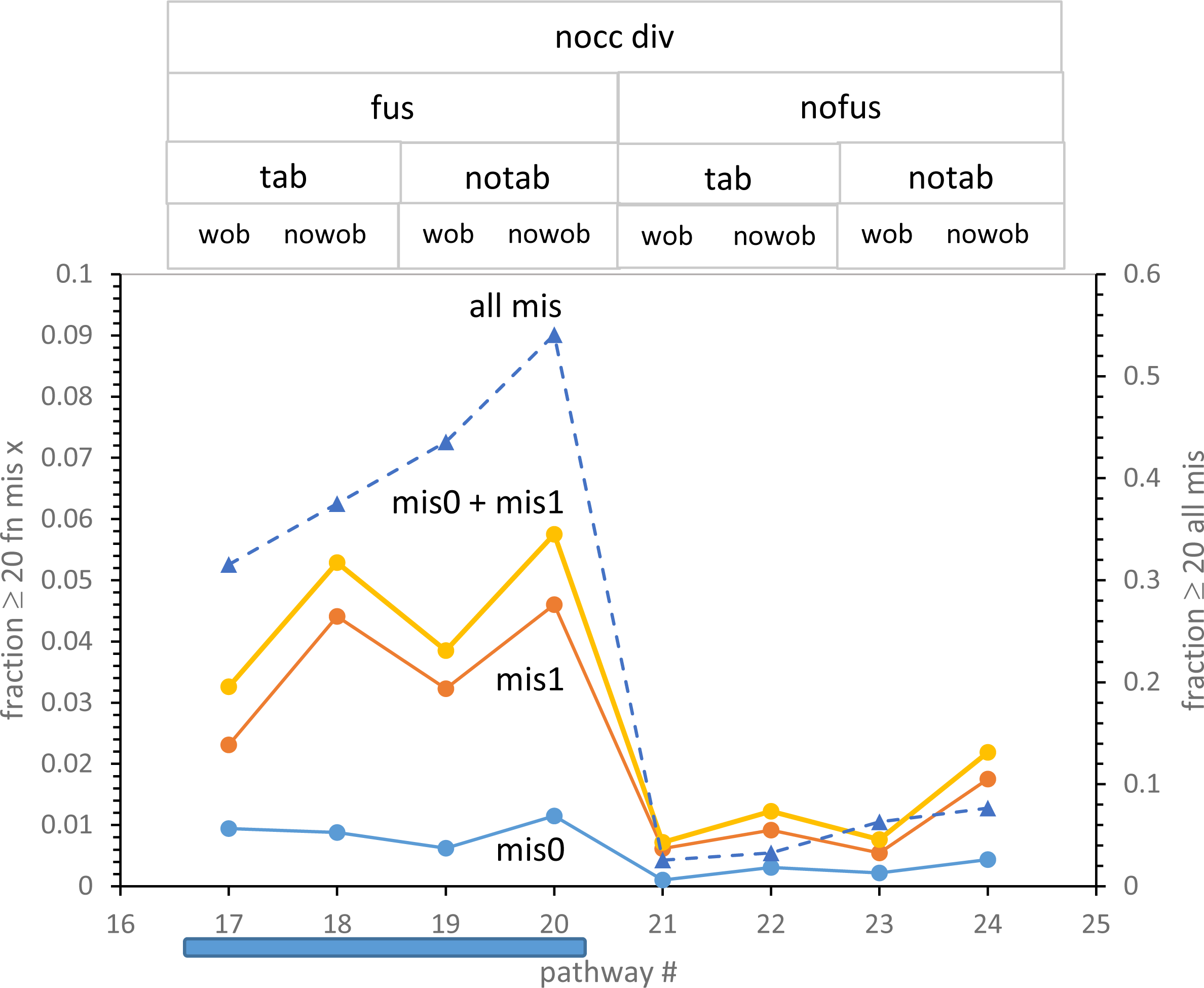
Fraction of codes with both ≥ 20 encoded functions and mis0 or mis1, or ≥ 20 encoded functions with any mis. All environments have run for 121 passages, the mean time for pathway #20 to reach ≥ 20 encoded functions. The shaded bar beneath the x-axis marks the superior nocc div fus section of the nocc div canyon pathways. Conditions are those of Fig. 1 except: Ptab = 0.08 or 0.0, Pwob = 0.005 or 0.0. Fractions are mean proportions of 1000 environments.

### Least selection resolves a fusion effect on accuracy

Fig. 8’s refined distance index resolves canyon pathways. There is a rift in the canyon floor: leftward pathways in Fig. 8 are much closer to the SGC than rightward. Consulting topward abbreviations, fusing pathways (#17-20) are much closer to the SGC than non-fusing ones (#21-24). Such evolutions may also employ independent tables or not (tab/notab), and/or may use wobble assignments or not (wob/nowob), but fusing routes remain always closer to the SGC. This resembles prior conclusions (Yarus 2021c, 2022a) that identified code fusions as decisive for rapid construction of SGC-like codes.

A canyon mechanism worth noting is #24, which relies on non-selective code division alone, nocc div nofus notab nowob. It is significantly slower than #18 and #20 to complete codes (Fig. 4) but has very good overall error (Fig. 6), and is deficient only in total SGC proximity (Fig. 8). Code division, even acting alone in pathway #24, suffices for moderately rapid code evolution.

Further, Fig. 8 again favors the exclusion of wobble during assignment (Yarus 2021b; as in Fig. 4 and Fig. 6) and also favors absence of parallel independent codes (notab) in the two mechanistic environments where it can be compared with a similar path (#17 vs #19 and also #18 vs #20).

### Four most competent pathways

Thus the favored path to the SGC is via the leftward canyon, nocc div fus. Moreover, the most favored pathway is resolved. That is #20, nocc div fus notab nowob.

But, given that choice, tab/notab and wob/nowob options are similar (differing by << 2-fold). At the early times of Fig. 8, for example, near-complete codes identical to the SGC using the second-best pathway #18 are 77% the abundance of similar codes via pathway #20. Thus, as a potential SGC pathway, #18 must be considered.

### Code division and fusion collaborate, but independently

It is no surprise that among the most SGC-like codes here, code fusion is frequent. Probabilities were chosen to make fusion effective. But a new question arises from the introduction of code division. Are division and fusion related, or independent features of code evolution? Though no div fus interaction was consciously implemented, human intuition is untrustworthy when so many processes interact.

Fig. 9 plots the product of the fraction of best codes fusing with the fraction dividing, for the 8/32 pathways that use both div and fus, and the 24 that do not (plotted at zero). This is compared to observation: the fraction of best codes employing both fusion and division are counted among results. Fig. 9 shows that the product of fraction fus and div and observed conjoined fusdiv in results are virtually identical. Therefore fusion and newly-introduced division aid code evolution, but by acting independently.

**Figure 9.**
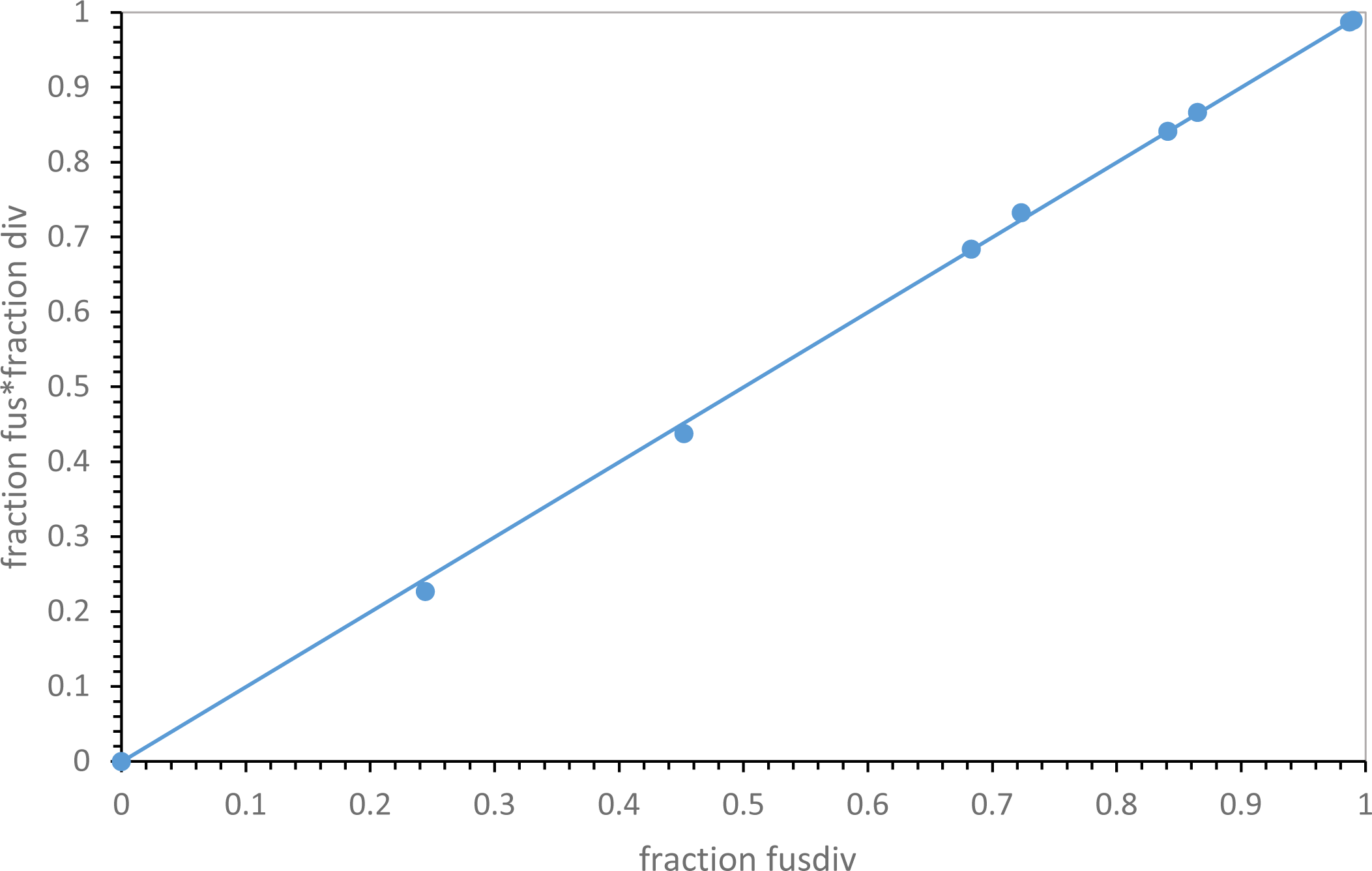
Independence of code fusion and division. Fraction of observed codes using fusion times fraction of observed codes using division versus fraction observed codes using division and fusion together. Environments are those in Fig. 4.

### A second, more efficient crescendo

Fig. 10 shows early kinetics for the reproducibly superior #20 pathway. In particular, it shows two code species closest to the SGC (≥ 20 encoded functions with mis0 or mis1). There is a rapid rise, then a prolonged presence of ≥ 20 assignment codes, no or one assignment away from the SGC. This accurate era lasts hundreds of passages. Thus, there will be many code assignments, decays, captures, fusions and divisions during this period. Said another way, proficient Fig. 10 codes will vary across time, but continuously present novel near-SGC-relatives for selection.

**Figure 10.**
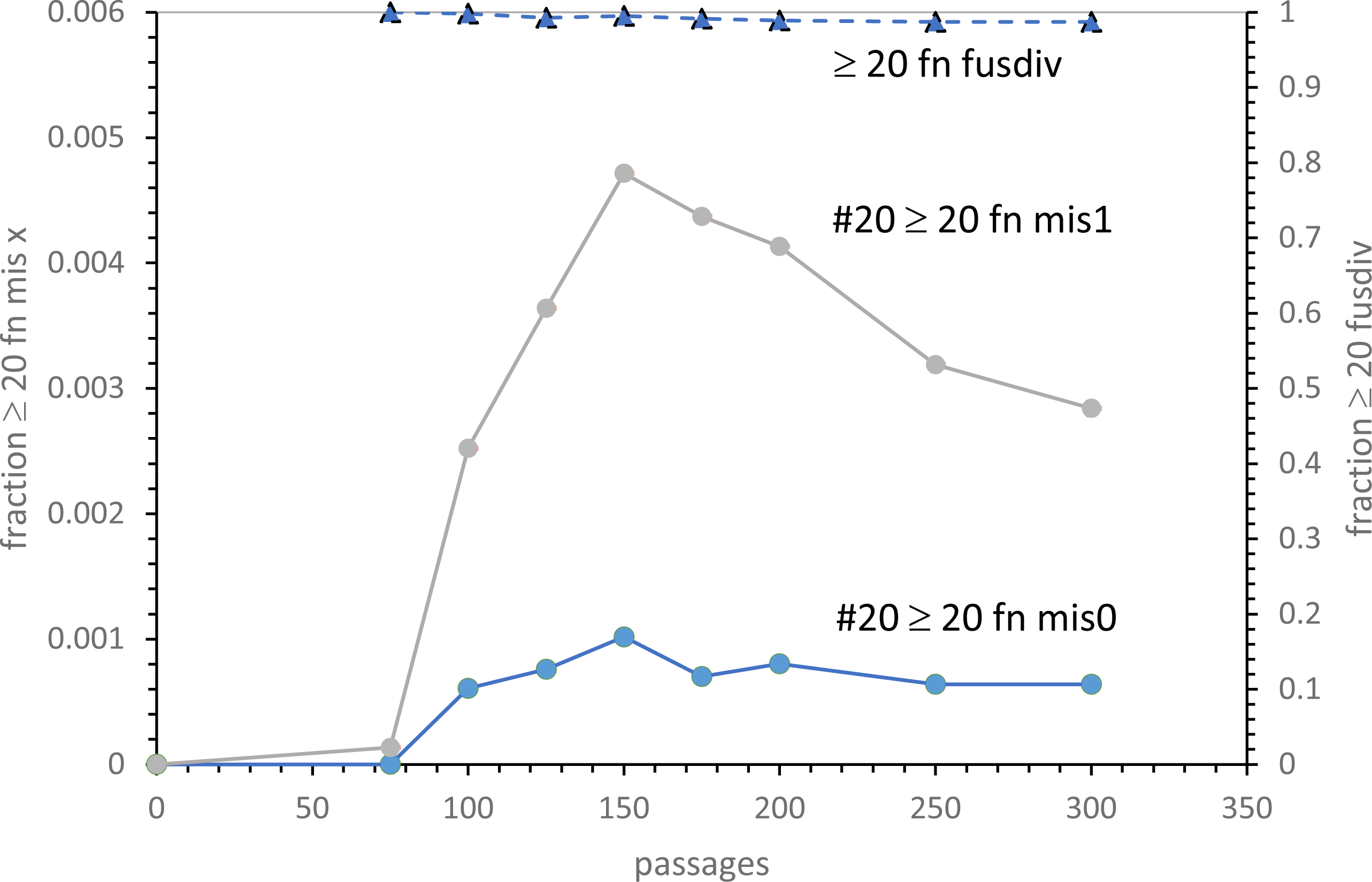
Kinetics of appearance of SGC-like codes. Codes with ≥ 20 encoded functions and also mis0 or mis1 are plotted versus time in passages. The fraction of ≥ 20 encoded function codes arising from fusion and division together (dashed line with triangles) is also plotted. Conditions are those for pathway #20: Pmut=0.00975, Pdec=0.00975, Pinit=0.15, Prand=0.05, Pfus=0.001, Ptab=0.0, Pwob=0.0, Pdivn=0.25, cc ≥ 1 encoded function.

Moreover, at Fig. 10 top is the fraction of ≥ 20 function codes that have assignments from both code fusion and division. Best SGC candidates arise nearly entirely by code division and fusion combined (dashed line, Fig. 10). This parallels the succession of highly competent codes from fus alone (Yarus 2022a), but this fus div crescendo arises more quickly and yields more frequent SGC-like codes. At the 150 passage peak there are 1 in 340 live ≥ 20 function mis0 codes (1 in 1000 total codes, including unsuccessful fusions: Fig. 10), or 1 in 74 live ≥ 20 function mis1 codes (1/210 of total codes: Fig. 10). Selection seems capable of finding these SGC-like codes. Fusion with division is a more probable route to the SGC than fusion alone (Yarus 2022b).

## Discussion

### Code division is influential

In this work, early genetic codes divide to make replicas of themselves. Code divisions are controlled by a probability of division per passage (Pdiv), and independently by a coding threshold (cc) that must be equaled or exceeded to make code division possible. The threshold is inspired by studies showing that 10 – 13 amino acids must be used to make active enzymes, and thus presumably, to make sophisticated structural proteins needed to support precise division. However, by reducing the threshold to 1 encoded amino acid, the threshold effect is circumvented – any existing code can then divide. It was thought to study the idea (Yarus 2022b) that the early code insured its evolutionary success by enabling its carrier to divide accurately; an evolutionary radiation whose winners had uniquely efficient protein biosynthesis then carried their genetic code to predominance.

Such an era has already been persuasively modeled (Vetsigian et al. 2006). Though late code-based radiation is still probable, results here concern an earlier era - code division profoundly alters nascent code history. Division speeds SGC evolution itself (Fig. 1A, 1B). Fastest evolution occurs for unselective division, when any code can divide (Fig. 1A, 3, 5). Under these conditions, the fastest approach to the SGC yet seen is observed (compare Yarus 2022a). Moreover, code division reinforces majority SGC assignments: when a mixture of SGC and random assignments is supplied, division tends to SGC rather than random assignments (Fig. 3). Preservation increases if division is more likely (increased Pdiv, Fig. 1B), as well as with more time to divide (cc = 1: Fig. 1A).

### Evolution in parallel

It was initially thought that code fusion would be advantageous because it allows parallel progress toward the SGC, gathering change made in different coding compartments instead of waiting for all modifications in a single ancestral line (Yarus 2021c). This can be quantitated (Fig. 4, 8; (Yarus 2022a). Here, in part because of improved fusion, dividing SGC evolution is ≈ 3-fold accelerated over non-dividing codes fusing with independent genetic codes (Fig. 4, 8, 9).

### 32 possible pathways to the SGC: rates of evolution

With addition of options for a code division threshold (cc/nocc) and division frequency (Pdiv ≥ 0) to previous models, there are 2^5^ = 32 different pathways for code evolution. Here, a code evolves entirely without one of these five effects, or in contrast, with a probability known to alter coding outcomes (see the supplementary data file).

Plotting evolutionary results against a structured list of pathways (Fig. 1A, 3, 4, 6, 12), defined at plot top, multiple different evolutionary pathways can be compared. This is first used for differing division rates and differing thresholds (Fig. 1A, 3) and then extended to the 32 pathways (Fig. 4, 6), emphasizing the rate of approach to the complete set of SGC assignments (Fig. 4), the adherence of the resulting codes to SGC encoding (Fig. 3, 6, 8) and the role of code division (Fig. 11).

**Figure 11.**
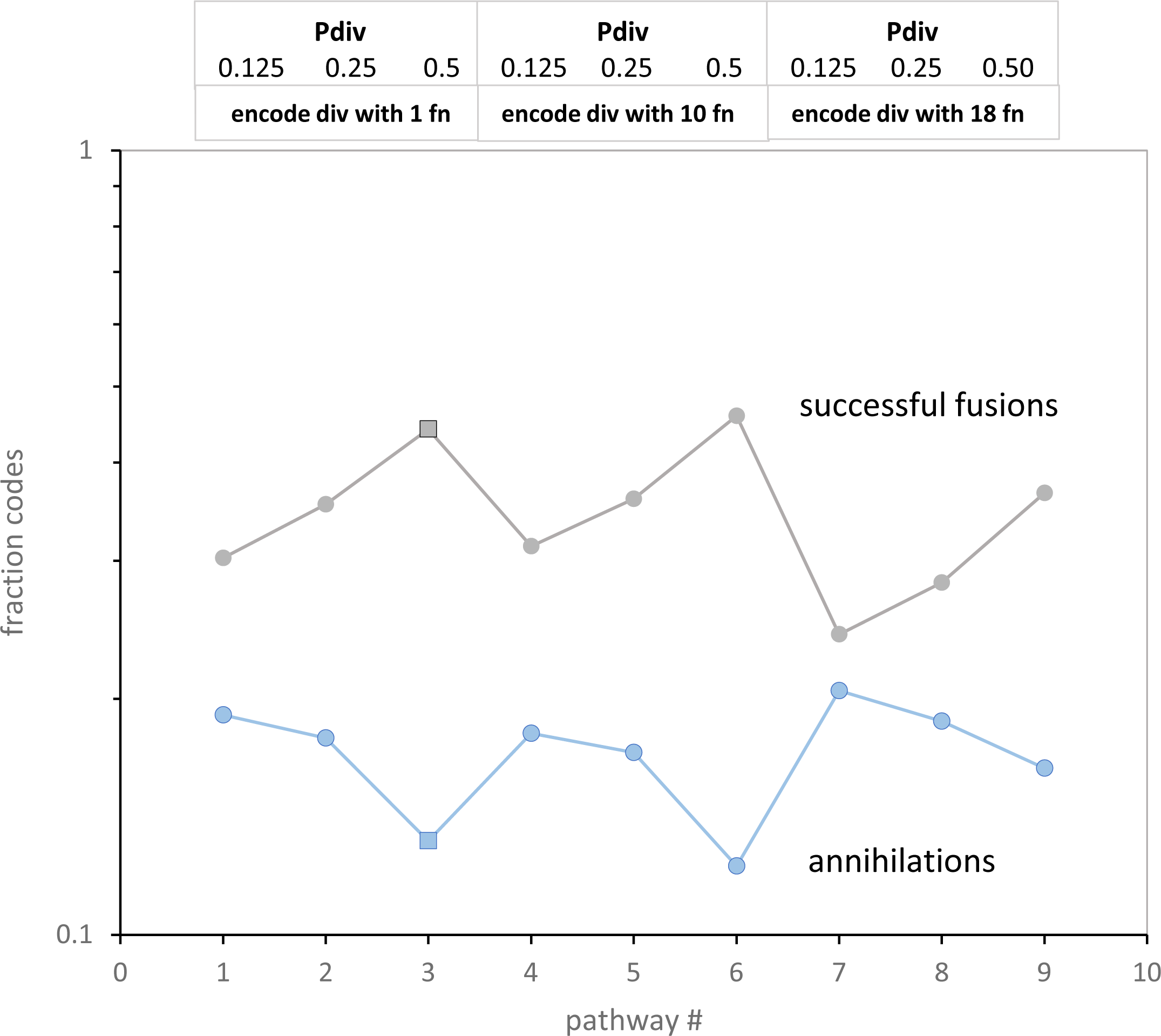
Successful and unsuccessful fusions (annihilations) have complementary behavior. Only code division varies; pathways and conditions are those of Fig. 1A, as indicated by abbreviations at plot top. Square points are the same as in Fig. 1A, 1B and 3.

The rate of evolution shows a notable canyon of fast rates (Fig. 4) for 8 mechanisms (#17-24) that allow code division (div) and impose no threshold for division (nocc). Conspicuously, all 8 canyon mechanisms encode ≥ 20 functions more quickly than any of the other 24 possible pathways.

### 32 possible pathways to the SGC: accuracy of evolution

There is a general relation between speed and accuracy of code evolution: this is shown in Fig. 7, where times for evolution to ≥ 20 functions is shown versus accompanying misassignment for 32 pathways. The observation that much of the variance for accuracy can be explained by rate of evolution is welcome. Fig. 7 implies one can find quick evolution accompanied by accurate assignment, so starting from a mixture of initial encodings becomes plausible.

This promise is fulfilled in Fig. 4 for rates and Fig. 6 for accuracies. These profiles have similar shapes: time to ≥ 20 functions and misassignments track well for most of 32 very different mechanisms (Fig. 7). Most particularly, a #17-24 canyon with quick evolution and accurate assignments exists in Fig. 4 and 6.

### A rift in the canyon floor: the role of fus

Codes requiring least selection (Yarus 2022b) to become the SGC, are likely precursors to the historical code. Thus, further resolution comes from a more precise measure of distance to the SGC, incorporating both speed and accuracy. In Fig. 8 this is implemented for the eight canyon codes (Fig. 4, 7), using as distance metric the fraction of codes that encode ≥ 20 functions and are also simultaneously accurate: mis0, mis1 or their sum.

Reading the upper legend of Fig. 8, there is a large difference in codes that fuse (fus) and those that do not (nofus). Maximally complete codes via fusion (#17-20) are about an order of magnitude more abundant than via non-fusing pathways (#21-24). This parallels previous findings (Yarus 2022a, 2023) that most complete codes come from code fusion. A nocc div fus canyon subset (Fig. 8) of pathways implement the doubly-capable evolution implied by the speed-accuracy correlation (Fig. 7).

Moreover, common use of code fusion by #17-20, the four most probable of 32 SGC pathways, supports the necessity of merging primordial codes, initially proposed for other reasons (Yarus 2021c, 2022a).

While differences among Fig. 8’s complete and accurate codes are not large, pathway #20 (nocc div fus notab nowob) is again superior, implying least selection to evolve the SGC.

### Implications of a flat canyon floor

Differences between tab/notab and wob/nowob codes are dramatically curtailed within the nocc div canyon, where these variations have their smallest observed effects (Fig. 4, 7). Such small effects are of evolutionary importance in two ways.

The first relates to wobble: how can one rationalize universal adoption of wobble coding when it is everywhere unfavorable (see **Wobble is always inhibitory** above)? One response is that wobble is likely delayed (Yarus 2021c), but another is that there exist pathways (#17-20, Fig. 4, 7, 9) where wobble has minimal effect. Wobble introduced late in pathway #18 or 20 would not be selected against.

### Comparing the best pathways

Small canyon-floor differences between tab/notab pathways are also evolutionarily significant. Routes #18 and 20 host codes that reach the SGC most quickly (Fig. 4) while also preserving high accuracy (Fig. 6). When speed and accuracy are required together (Fig. 8), #18 and 20 are again best. What does this multiple superiority mean?

Fig. 8 shows that #18 and #20 environments differ only for independent codes - it is somewhat better to avoid them. This is puzzling, because more independent codes provide a broader sample of the coding environment, and are generally expected to find the SGC sooner (Yarus 2022a). Moreover, multiple codes can fuse, quickly forming more complete codes by summing compatible assignments (Yarus 2021c, 2022a). Fig. 8 therefore suggests that something subtle makes path #20 (nocc div fus notab nowob) best, and in particular, superior to #18 (nocc div fus tab nowob).

### Multiple codes are more advantageous if they resemble each other

The subtlety is in the nature of “other” codes. When independent codes fuse, they do assemble complete, accurate code tables significantly more rapidly (Yarus 2022a). But code division creates a new kind of fusion partner. Fig. 11 shows this, using a pathway using both independent and division partners (Fig. 1A). As code division increases in Fig. 11, the fraction of codes with successful fusion among environmental codes increases.

Even more relevantly, unsuccessful fusions (annihilations via conflicting assignment) decrease, and by the same proportion as fusion increase. Fig. 11’s two plots mirror each other. Especially apt fusion partners from code division replace fusion to independently arising codes, to make up the approximately constant number of assignments required for complete code construction (Fig. 1A). However, at all levels of code division, quickest (Fig. 1A) and most accurate SGC-like evolution (Fig. 3) is associated with the greatest successful (Fig. 11, square) and least unsuccessful (Fig. 11) code fusion.

With time, dividing, highly related, code numbers increase, so variants of a dividing code will be made and tested more rapidly. This resembles the ‘crescendo’ of competent codes created by fusion with increasing numbers of unrelated codes (Yarus 2022a). In this work, novel partner codes originate by division and subsequent evolution, but the result is similar: an era when highly complete, highly accurate codes proliferate. SGC selection can survey a second kind of prolonged fusion-division crescendo (Fig. 11), during which many different-but-related SGC-like codes can be assessed.

### Pathway #20 simplifies SGC evolution

Thus the “disadvantage” of fusion between independent codes is only that a better path exists: a dividing code population harvests evolutionary change by fusing evolved ancestral codes and varied descendants. Most especially, this effect makes evolution from a unique origin (pathway #20) even <u>more</u> efficient than fusing with independent codes (pathway #18). A simpler primordial code emergence by least SGC selection from a single ancestor becomes plausible.

### Hybrid routes to the SGC

Fig. 1’s varying division rates move code evolution along an axis joining pathway #18 and #20. Code division increases, independent code fusion decreases (toward #20), or the reverse (toward #18) with small change in result (Fig. 8). Hybrid routes with similar SGC access suggest novel possibilities. Specifically, pathways #18 and 20 approach the SGC by fusing dissimilar early coding tables. Therefore, they suggest that other partial codes could be fused.

The SGC can have an even earlier history (Fontecilla-Camps 2023), but the early code is usually structured in one of four ways. ‘Frozen accidents’: (Crick 1968) supposed that a code could be frozen, perhaps after being shaped by earlier molecular interactions. In any case, a growing code would become difficult to change because changes would perturb all previous gene products (Sella and Ardell 2006). ‘Coevolution’: as (Koonin and Novozhilov 2008) have emphasized, it is undeniable that code progress could have been shaped by metabolic evolution, more complicated amino acids encoded only after progressive biosynthesis reaches them. This is a highly developed theory (Wong 1975; Taylor and Coates 1989; Di Giulio 2008; Higgs and Pudritz 2009) often called coevolution of the genetic code. ‘Error minimization’: a code or partial code might be shaped by selection to minimize the effects of coding errors or mutations (Haig and Hurst 1991; Freeland and Hurst 1998). Strikingly, error minimization can arise without selection against error (Massey 2016). ‘Stereochemistry’: coding assignments might reflect chemical interaction of amino acids and ribonucleotides. Selected RNA binding sites for amino acids contain an excess of anticodon and codon triplets. Each triplet is an essential binding sequence, as shown by sequence conservation and mutagenesis data (Yarus et al. 2005; Rodin et al. 2011; Yarus 2017). Genomic sequencing (Bartonek and Zagrovic 2017; Kapral et al. 2022) suggests that related interactions are recorded throughout modern mRNAs.

Notably, all mechanisms could yield code fragments that fuse. Even more to this point, divergent mechanisms plausibly utilize varied sets of triplets. Codes from disparate origins could have fewer overlapping, conflicting assignments. As for independent codes versus dividing codes (Fig. 8, 11), efficient evolution results when code fusions are less failure-prone.

### Biology as anthology

Inspired by calculated advantages of code fusion, it was suggested that life can be defined by facile gathering of separate advantages into one line of descent (Yarus 2022b). From this work, we add that life can automatically refine gathered advantages (Fig. 4, 7), but also effortlessly combine advantages (Fig. 11). Division and fusion are elementary cellular activities; thus code evolution has a simple, almost inevitable, rationale. Such powerful cellular effects were probably not used solely to create the SGC.

## Methods

### Monte Carlo kinetics

Imagine recording events by quickly making a mark on a chart moving past you across a table: for coding evolution, these marks represent codon assignments, captures, code divisions, and so on. The table is not passive: it both records marks and allows them to evolve according to physical and chemical laws. At the end of evolution, lay a laddered grid over your series of chart marks. The ladder has narrow gaps; only one event (or no event) ever appears in an inter-step opening of your ladder. In placing your grid, you have not changed the sequence of events and notably, not changed the evolutionary result on the table. Thus, you have shown that you can reproduce a “continuous” kinetic process with small adjacent windows (here called passages), assigning probabilities for each kind of event within a window. If the probability of a mark on the chart depends on the concentration of something on the table, you are modeling 1^st^-order kinetics, and defining a first order rate constant (Yarus 2021b). If probability of appearing in a window/passage depends on two concentrations, you are modeling 2^nd^-order kinetics. This Monte Carlo framework makes it simple to implement many rates you wish to study, even in complex environments like that on your (coding) table, with many changes occurring at many loci for change. Resulting source code in Pascal is available on request from the author. An array of 32 named pathways with selected results is in a supplementary manuscript file.

### Tracking time

Time passes for environmental codes in passages. During a passage, every existing coding table is given the chance to <u>either</u> assign a randomly-chosen, unassigned codon (probability Pinit), have a previous assignment decay (Pdecay), or capture an unassigned codon a single mutation distant (Pmut), conferring on it an assignment with a closely related polar requirement (Woese 1965; Mathew and Luthey-Schulten 2008), including transferring the assignment of the initial codon. The final options are that before internal passages, two coding tables may fuse (Pfus) – this can augment code growth if the fusees combine compatible assignments, or it can inactivate fusees if their pre-existing codes make differing assignments to the same codon. Incompatible codes are inactivated, and do not evolve further (Yarus 2022a). Alternatively, an existing code accurately divides (Pdiv).

### Coding and wobble coding

Functions are encoded by assignment to a single coding triplet, or via simplified Crick wobble, which allows U/G and G/U pairs at the third position of codons (Crick 1966; Yarus 2021b). Onset of wobble is controlled by Pwob, the probability of wobble onset per passage (Yarus 2023).

## Supporting information

Structured list and data for 32 pathways

Excel list and data for 32 pathways

