## Supplementary material for "The genetic code assembles via division and fusion, basic cellular events": Structured list and data for 32 pathways

supplementary summary of selected output sorted by pathway #

| | | constants | | | | | results | | | | | | | | | | |
|---|---|---|---|---|---|---|---|---|---|---|---|---|---|---|---|---|---|
| **pathway abbreviations** | ***pathway #*** | **cc** | **div** | **fus** | **tab** | **wob** | **envi** | **tadx** | **pgdx** | **inipsg** | **Fus** | **mis_W** | **mis** | **nrdiv** | **livtab** | **fr MxC >=20, mis0** | **fr MxC >=20, mis1** |
| cc div fus tab wob | ***1*** | 0.81 | 0.25 | 0.001 | 0.08 | 0.005 | 1000 | 56.9348 | 442.5055 | 231.0602 | 23.10933 | 7.148445 | 6.224674 | 2.872618 | 15.00702 | 0.01003009 | 0.033099298 |
| cc div fus tab nowob | ***2*** | 0.81 | 0.25 | 0.001 | 0.08 | 0 | 1000 | 22.60262 | 255.9849 | 119.339 | 20.92052 | 6.257545 | 4.118712 | 1.652918 | | 0.024144869 | 0.072434608 |
| cc div fus notab wob | ***3*** | 0.81 | 0.25 | 0.001 | 0 | 0.005 | 500 | 83.208 | 1421.678 | 0 | 3.744 | 11.262 | 10.414 | 8.62 | 14.86 | 0 | 0.016 |
| cc div fus notab nowob | ***4*** | 0.81 | 0.25 | 0.001 | 0 | 0 | 500 | 15.1984 | 434.2084 | 0 | 0.945892 | 9.412826 | 7.086172 | 3.128257 | 11.53307 | 0.002004008 | 0.022044088 |
| cc div nofus tab wob | ***5*** | 0.81 | 0.25 | 0 | 0.08 | 0.005 | 1000 | 27.54545 | 314.9151 | 36.04196 | 0 | 7.966034 | 6.771229 | 2.12987 | 38.04196 | 0.008991009 | 0.026973027 |
| cc div nofus tab nowob | ***6*** | 0.81 | 0.25 | 0 | 0.08 | 0 | 1000 | 23.06593 | 280.0899 | 31.12887 | 0 | 7.256743 | 4.647353 | 1.816184 | 31.37962 | 0.013986014 | 0.03996004 |
| cc div nofus notab wob | ***7*** | 0.81 | 0.25 | 0 | 0 | 0.005 | 499 | 22.76553 | 1054.168 | 0 | 0 | 10.95992 | 10.1002 | 3.731463 | 35.11623 | 0.002004008 | 0.01002004 |
| cc div nofus notab nowob | ***8*** | 0.81 | 0.25 | 0 | 0 | 0 | 500 | 8.88 | 391.508 | 0 | 0 | 9.172 | 6.628 | 2.66 | 14.48 | 0.004 | 0.012 |
| cc nodiv fus tab wob | ***9*** | 0.81 | 0 | 0.001 | 0.08 | 0.005 | 250 | 124.124 | 1713.732 | 1533.184 | 24.144 | 6.896 | 6.336 | 0 | 10.036 | 0.012 | 0.048 |
| cc nodiv fus tab nowob | ***10*** | 0.81 | 0 | 0.001 | 0.08 | 0 | 250 | 23.456 | 446.996 | 278.676 | 20.28 | 6.736 | 4.608 | 0 | 10.488 | 0.016 | 0.088 |
| cc nodiv fus notab wob | ***11*** | 0.81 | 0 | 0.001 | 0 | 0.005 | 250 | 1 | 29696.61 | 0 | 0 | 14.74 | 14.236 | 0 | 1 | 0.004 | 0 |
| cc nodiv fus notab nowob | ***12*** | 0.81 | 0 | 0.001 | 0 | 0 | 250 | 1 | 1344.844 | 0 | 0 | 11.568 | 9.676 | 0 | 1 | 0 | 0.004 |
| cc nodiv nofus tab wob | ***13*** | 0.81 | 0 | 0 | 0.08 | 0.005 | 250 | 7.702811 | 423.4699 | 81.53414 | 0 | 8.566265 | 7.570281 | 0 | 34.75502 | 0 | 0.020080321 |
| cc nodiv nofus tab nowob | ***14*** | 0.81 | 0 | 0 | 0.08 | 0 | 250 | 5.64 | 349.748 | 58.396 | 0 | 7.968 | 5.3 | 0 | 28.56 | 0.008 | 0.02 |
| cc nodiv nofus notab wob | ***15*** | 0.81 | 0 | 0 | 0 | 0.005 | 250 | 1 | 18031.55 | 0 | 0 | 14.136 | 13.572 | 0 | 1 | 0 | 0.004 |
| cc nodiv nofus notab nowob | ***16*** | 0.81 | 0 | 0 | 0 | 0 | 250 | 1 | 1586.136 | 0 | 0 | 11.712 | 9.904 | 0 | 1 | 0.008 | 0.024 |
| nocc div fus tab wob | ***17*** | 0.045 | 0.25 | 0.001 | 0.08 | 0.005 | 1000 | 37.58042 | 177.5455 | 45.35165 | 23.1978 | 6.668332 | 5.10989 | 4.195804 | | 0.016983017 | 0.065934066 |
| nocc div fus tab nowob | ***18*** | 0.045 | 0.25 | 0.001 | 0.08 | 0 | 1000 | 25.902 | 132.518 | 22.904 | 22.013 | 5.955 | 3.859 | 3.542 | | 0.029 | 0.099 |
| nocc div fus notab wob | ***19*** | 0.045 | 0.25 | 0.001 | 0 | 0.005 | 999 | 34.78773 | 165.9567 | 0 | 18.42857 | 6.720322 | 5.150905 | 7.158954 | 16.78068 | 0.019114688 | 0.069416499 |
| nocc div fus notab nowob | ***20*** | 0.045 | 0.25 | 0.001 | 0 | 0 | 1000 | 23.825 | 121.331 | 0 | 17.629 | 6.112 | 3.894 | 5.657 | | 0.024 | 0.089 |
| nocc div nofus tab wob | ***21*** | 0.045 | 0.25 | 0 | 0.08 | 0.005 | 1000 | 38.35276 | 185.5116 | 2.043216 | 0 | 6.650251 | 5.073367 | 6.744724 | 47.22513 | 0.020100503 | 0.060301508 |
| nocc div nofus tab nowob | ***22*** | 0.045 | 0.25 | 0 | 0.08 | 0 | 1000 | 36.94494 | 174.6106 | 1.564565 | 0 | 6.006006 | 3.55956 | 6.825826 | | 0.043043043 | 0.105105105 |
| nocc div nofus notab wob | ***23*** | 0.045 | 0.25 | 0 | 0 | 0.005 | 1000 | 35.24525 | 164.1111 | 0 | 0 | 6.06006 | 4.437437 | 8.850851 | 41.63363 | 0.031031031 | 0.093093093 |
| nocc div nofus notab nowob | ***24*** | 0.045 | 0.25 | 0 | 0 | 0 | 1000 | 34.34036 | 157.0843 | 0 | 0 | 5.797189 | 3.349398 | 8.720884 | 40.16265 | 0.039156627 | 0.124497992 |
| nocc nodiv fus tab wob | ***25*** | 0.045 | 0 | 0.001 | 0.08 | 0.005 | 250 | 113.12 | 1563.396 | 1390.712 | 24.048 | 6.86 | 6.284 | 0 | 10.132 | 0.008 | 0.032 |
| nocc nodiv fus tab nowob | ***26*** | 0.045 | 0 | 0.001 | 0.08 | 0 | 1000 | 24.75876 | 433.6056 | 290.3133 | 21.60561 | 6.276276 | 4.237237 | 0 | | 0.021021021 | 0.066066066 |
| nocc nodiv fus notab wob | ***27*** | 0.045 | 0 | 0.001 | 0 | 0.005 | 250 | 1 | 31059.34 | 0 | 0 | 15.22 | 14.756 | 0 | 1 | 0 | 0 |
| nocc nodiv fus notab nowob | ***28*** | 0.045 | 0 | 0.001 | 0 | 0 | 250 | 1 | 1427.452 | 0 | 0 | 12.392 | 10.384 | 0 | 1 | 0 | 0 |
| nocc nodiv nofus tab wob | ***29*** | 0.045 | 0 | 0 | 0.08 | 0.005 | 250 | 7.332 | 419.892 | 80.172 | 0 | 8.952 | 8.1 | 0 | 34.384 | 0.008 | 0.008 |
| nocc nodiv nofus tab nowob | ***30*** | 0.045 | 0 | 0 | 0.08 | 0 | 250 | 5.257028 | 339.1767 | 47.88755 | 0 | 7.614458 | 5.076305 | 0 | 27.95181 | 0.016064257 | 0.02811245 |
| nocc nodiv nofus notab wob | ***31*** | 0.045 | 0 | 0 | 0 | 0.005 | 250 | 1 | 15403.51 | 0 | 0 | 14.288 | 13.756 | 0 | 1 | 0 | 0 |
| nocc nodiv nofus notab nowob | ***32*** | 0.045 | 0 | 0 | 0 | 0 | 250 | 1 | 1568.364 | 0 | 0 | 12.34 | 10.708 | 0 | 1 | 0 | 0.004 |
| | | **cc** | **div** | **fus** | **tab** | **wob** | **envi** | **tadx** | **pgdx** | **inipsg** | **Fus** | **mis_W** | **mis** | **nrdiv** | **livtab** | **fr MxC >=20, mis0** | **fr MxC >=20, mis1** |

***key for pathway constants***

**cc =** completeness criterion, slightly < (assignments needed to support code division/22)

**div** = Pdiv, probability of code division during a passage

**fus** = Pfus, probability of fusion for a table during a passage

**tab** = Ptab, probability of addition of an independent code table during a passage

**wob** = Pwob, probabilty that wobble is initiated during a passage

All data are means for the tabulated number of environments.

All pathways have 5% random assignments.

**key for numerical results**

**envi**: number of environments

**tadx**: mean number of total coding tables

**pgdx**: number of passages @ ≥ 20 encoded functions

**inipsg**: passage at which ≥ 20 fn code was initiated

**Fus**: number of assignments by code fusion

**mis_W**: number of misassignments with completed simple Crick wobble

**mis**: number of initial misassignments relative to SGC

**nrdiv**: number of code divisions @ ≥ 20 functions

**livtab**: number of live coding tables @ ≥ 20 functions

**fr MxC >=20, mis0**: fract max complete codes ≥ 20 assigned fn, mis0

**fr MxC >=20, mis1**: fr max comp codes ≥ 20 fn, mis1
